# Using a robotic fish to investigate individual differences in social responsiveness in the guppy

**DOI:** 10.1101/304501

**Authors:** David Bierbach, Tim Landgraf, Pawel Romanczuk, Juliane Lukas, Hai Nguyen, Max Wolf, Jens Krause

## Abstract

Responding towards the actions of others is one of the most important behavioral traits whenever animals of the same species interact. Mutual influences among interacting individuals may modulate the social responsiveness seen and thus makes it often difficult to study the level and variation of individuality in responsiveness. Here, biomimetic robots (BRs) that are accepted as conspecifics but controlled by the experimenter can be a useful tool. Studying the interactions of live animals with BRs allows pinpointing the live animal’s level of responsiveness by removing confounding mutuality. In this paper, we show that live guppies (*Poecilia reticulata*) exhibit consistent differences among each other in their responsiveness when interacting with a biomimetic fish robot - ‘Robofish’ - and a live companion. It has been repeatedly suggested that social responsiveness correlates with other individual behavioral traits like risk-taking behavior (‘boldness’) or activity level. We tested this assumption in a second experiment. Interestingly, our detailed analysis of individual differences in social responsiveness using the Robofish, suggests that responsiveness is an independent trait, not part of a larger behavioral syndrome formed by boldness and activity.

## Introduction

Synchronized behaviors such as collective movements depend on the capability of involved subjects to respond to the actions of their social partners [1-6]. Such a responsiveness towards the social environment has been termed either ‘sociability’ ([7], the tendency to approach rather than to avoid conspecifics, see also [2, 8]), ‘social competence’ ([9], as an adaptive response in a social context) or ‘social responsiveness’ ([10], a tendency to respond to past or present reputation/action of conspecifics).

While there is some discussion regarding terminology (see [11]), assessing any response of an individual towards its social environment inevitably requires the presentation of social cues from conspecifics. The use of live conspecifics for this purpose typically is problematic as they often interact with the focal individual and thereby introduce confounding variation into the experimental design (see e.g. [12-15]). Thus, experimenters tried to control for or standardize the possible mutual interactions among subjects. Some studies used pretrained live “demonstrators” to interact with naïve individuals [16-21], while other approaches modified binary classical shoaling assays [22] in a way that individuals, although spatially separated, can decide whether to associate or not with a visible conspecific in an adjacent compartment [8, 23]. More recently, video playbacks or computer animations have been used to create and even manipulate social stimuli [24-31]. Similarly, others have presented live animals with spatially-separated artificial models of conspecifics [32-35]. However, realistic tests of social responsiveness towards movement patterns of social partners, as for example found in collectively moving shoals of fishes, herds of ungulates or flocking birds, require a spatial scale [36].

The need to provide a spatial scale while still be able to control for or standardize social interactions inspired experimenters to look for the interactions of live animals towards moving replicas (see [37-42]). Recently, sticklebacks have been found to differ consistently from each other in their attraction towards a dummy school that circulates at a constant speed [42], a technique that has been also used previously to investigate shoaling tendencies in blind cave tetras (*Astyanax mexicanus*, [40]). Even more sophisticated experiments became feasible through the development of biomimetic robots [36, 43, 44]. Biomimetic robots, that are accepted as conspecifics by live animals have several advantages over previous approaches, such as the ability to completely standardize the behavior of the interacting robot, to set its parameters to either resemble those of focal fish or show a sharp contrast with them, as well as to allow the possibility to create interactive scenarios that nevertheless follow controlled rules that can be adapted intentionally [45-48].

Using a biomimetic robot (hereafter called ‘Robofish’) that is accepted as a conspecific by live Trinidadian guppies (*Poecilia reticulata*; see [45]), we ask in the current study (a) whether animals differ in their social responsiveness and whether this difference can be measured with a biomimetic robot; (b) whether among-individual differences in social responsiveness towards moving (robotic) conspecifics are linked to other behavioral traits that have been established to differ among live animals (‘personality traits’ see [7]).

In our first experiment (question a), we specifically predicted that an individual’s level of responsiveness towards a live conspecific should resemble that towards moving Robofish. As our Robofish, although accepted as conspecific, is steered in an non-interactive open-loop mode to omit mutual influences on the live fish’s behavior, we do not expect that interactions with Robofish resemble fully that of live fish interactions but predict that interaction patterns should be similar. We measured several interaction parameters (inter-individual distance, velocity cross-correlations, Transfer Entropy) of focal fish with live partners and, in a subsequent test with Robofish partners and calculated behavioral repeatability, a measure for consistent individual differences [49]. If tests with Robofish are able to depict an individual’s responsiveness, among-individual differences should be consistently detectable also when the same focal fish is tested with a live partner.

It is known that behaviors often form correlated suits, so-called behavioral syndromes [50]. For example, it is known from studies on sticklebacks that individuals with increased tendencies to take risks (behavioral trait ‘boldness’ and/or ‘exploration behavior’) lead more often and are less attracted by others, while those with lower tendencies to take risks are more likely to follow others [8, 15]. As similar relations are possible in regard to the social responsiveness of an individual (question b), we predicted in our second experiment that focal fish that were highly responsive to Robofish’s actions should be less risk-taking and explorative while those that did not respond strongly to Robofish should be more risk taking and explorative.

## Methods

### Study organism and maintenance

We used wild-type guppies (*Poecilia reticulata*) for our experiments that have been bred in the laboratory for several generations and originated from wild-caught individuals. Test fish came from large, randomly outbred single-species stocks maintained at the animal care facilities at the Faculty of Life Sciences, Humboldt University of Berlin. We provided a natural 12:12h light:dark regime and maintained water temperature at 26°C. Fish were fed twice daily *ad libitum* with commercially available flake food (TetraMin™) and once a week with frozen *Artemia* shrimps.

### The Robofish system

The Robofish system consists of a glass tank (88 × 88 cm), which is mounted onto an aluminum rack. A two-wheeled robot can move freely on a transparent platform below the tank (figure 1A-B). The robot carries a magnet, coupling its motion with a second magnet in the tank above. The second magnet serves as the base for a three-dimensional 3D-printed fish replica (standard length (SL) =30.0 mm; resembling a guppy female, see figure 1C). This kind of replicas are accepted as conspecifics by live guppies (and other fishes), most likely through the use of glass eyes and by swimming in a natural motion pattern [45, 51]. The entire system is enclosed in a black, opaque canvas to minimize exposure to external disturbances. The tank is illuminated from above with artificial light reproducing the daylight spectrum. On the floor, a camera is facing upwards to track the robot. A second camera is fixed above the tank to track both live fish and the replica. Two computers are used for system operation: one PC tracks the robot, computes and sends motion commands to the unit over a wireless channel; the second PC records the video feed of the ceiling camera, which is subsequently tracked by a custom-made software [52].

**Figure 1:**
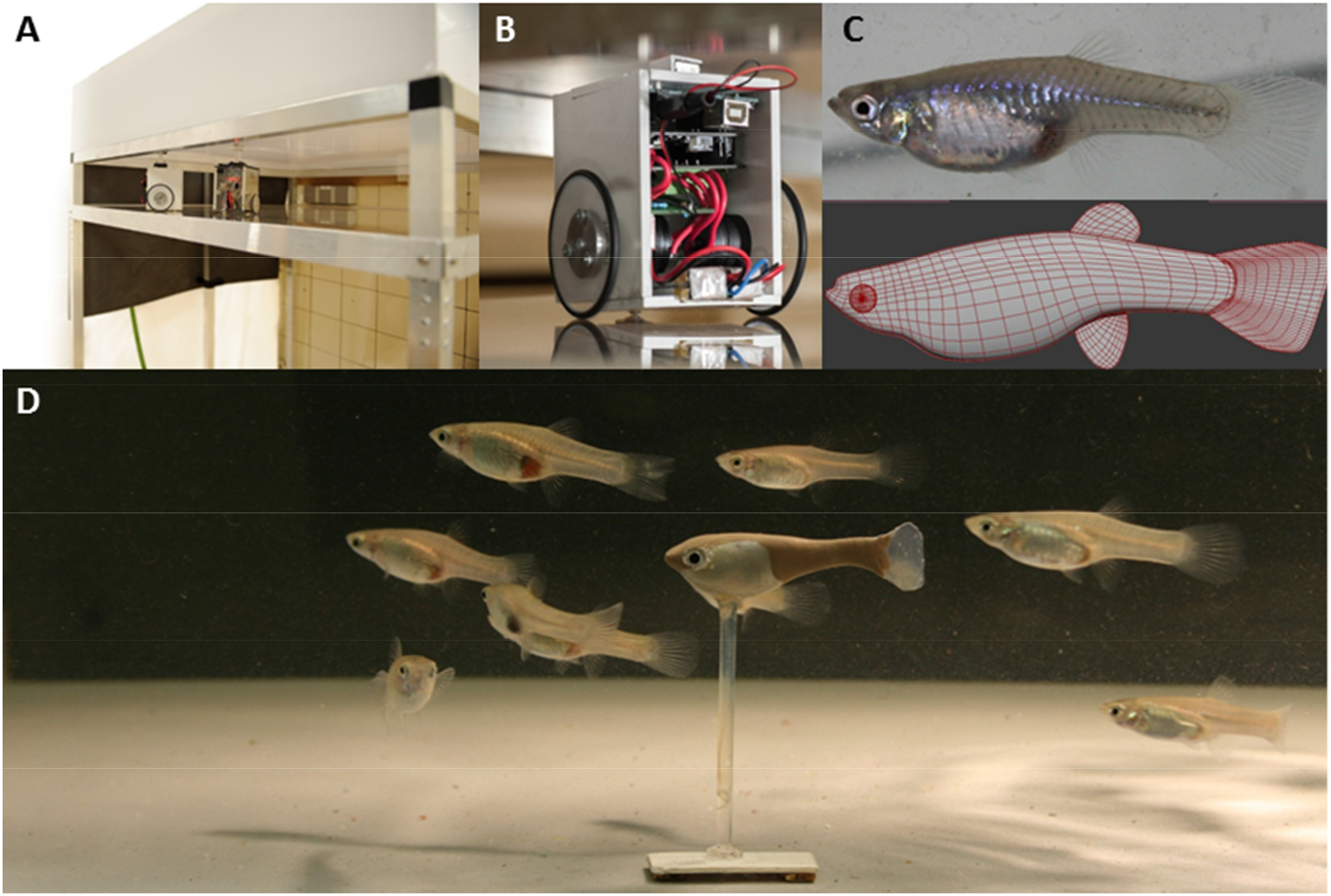
The Robofish system. (A) The robot unit is driving on a second level below the test arena. (B) Close-up of the robot unit. (C) A picture of a live guppy female served as template for the virtual 3D mesh that was printed on a 3D printer. (D) Guppy replica with a group of female guppies in the test arena.

#### Experiment 1

##### Experimental setup

To compare responses of live focal fish between tests with Robofish and with a live partner, each focal fish was tested once with Robofish and subsequently another time with a live model individual. This was done by testing one half of the focal fish first with Robofish and after two days with a live model fish, while the other half of the focal fish were first tested with a live model fish and after two days with Robofish. Focal fish were randomly assigned to start with the Robofish or live model treatment.

At the beginning of our experiment, we randomly selected adult fish from our stock tanks (females-only to reduce possible sex-specific differences) and marked them individually with VIE elastomeric color tags (see [53]). During this procedure, we also measured body length as standard length (from tip of snout to end of caudal peduncle) to the nearest millimeter (focal fish: SL ± SEM = 30.1 mm ± 0.4 mm, *N*=30; live model fish: 30.5 mm ± 0.3 mm, *N*=30).

To initiate a trial, we transferred half of the test fish (N=15) into a Plexiglas cylinder located at the upper left corner of the arena (see figure 2). The Robofish replica was also located within the cylinder. After a habituation period of 2 minutes, robot and live fish were released by lifting the cylinder with an automatic pulley system. When the live fish left the cylinder (= one body length away from the cylinder’s border), Robofish started swimming in a natural stop-and-go pattern [45, 54] along a zigzag path to the opposite corner (figure 2). After reaching this corner, the Robofish randomly swam to either the bottom left or the top right corner in which it ultimately described a circular path for three rounds (figure 2).

**Figure 2:**
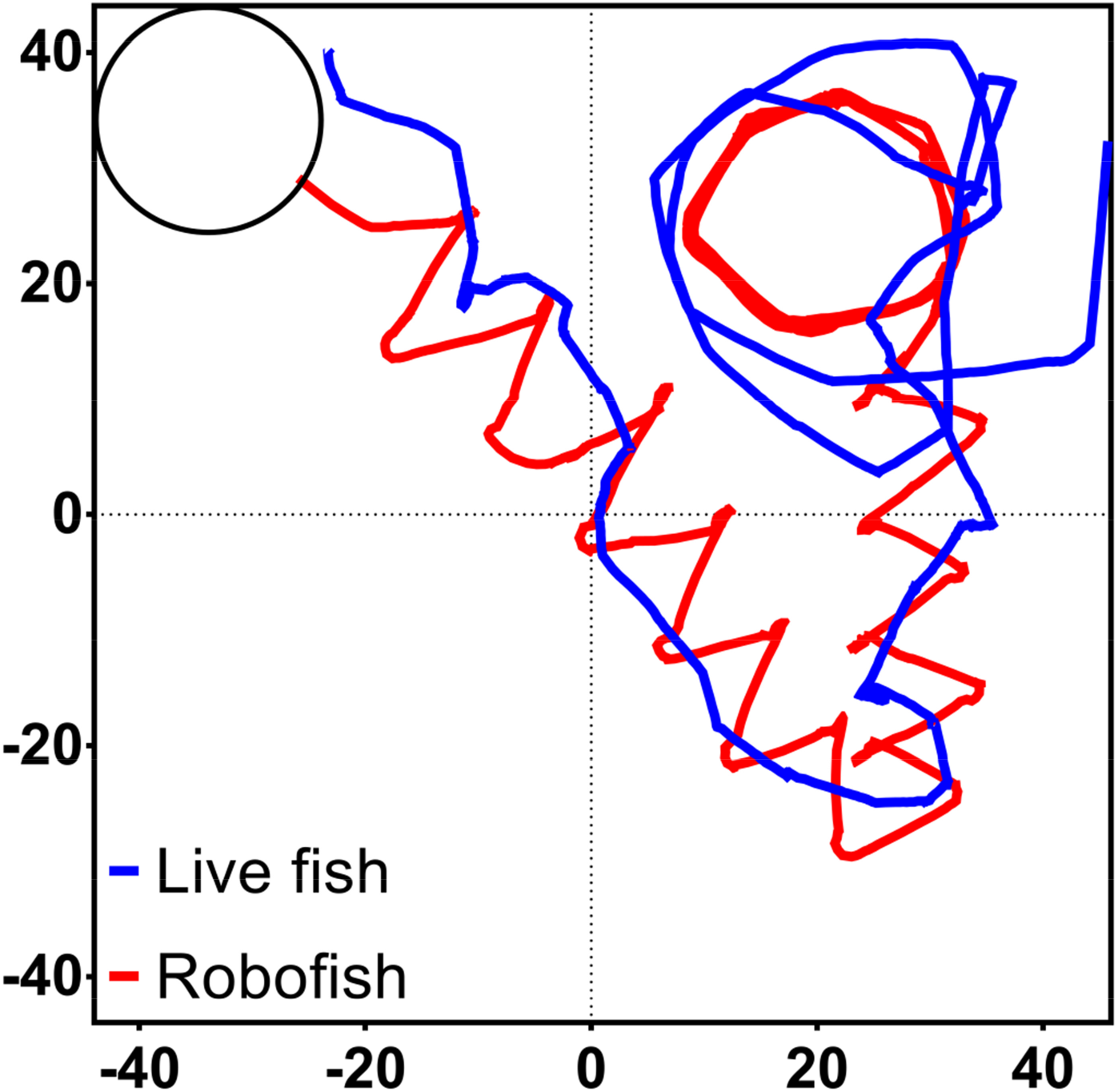
Example track of Robofish with live guppy in an 88cm x 88 cm test arena. After the live fish left the start cylinder (upper left), Robofish moved in a natural stop-and-go pattern along a zigzagged path to the opposite corner. Upon arrival, Robofish moved to either the bottom left or the top right corner (here: top right) and described a circular path.

The trial was then terminated and the test fish was transferred back to its holding tank. Each trial was videotaped for subsequent analysis. A video recording following this protocol is available as an online supplement (Video S1). To test whether the focal fish’s response towards Robofish can be linked to the response towards a live conspecific, the other half of the focal fish was instead introduced into the start cylinder accompanied with a live companion comparable in size to the Robofish replica (see above). Again, we lifted the cylinder after 2 min of habituation and videotaped the trial for 2 min, starting when the last fish left the cylinder. Trials involving only live fish were comparable in duration to Robofish trials (live-live: 120s; Robofish: 124.1 s ± 1.9 s; mean ± SEM, variation in duration is due to stop-and-go swimming pattern of Robofish). After two days the testing was repeated; however, fish were now introduced to the opposite treatment. Thus, each focal fish (N = 30) was tested with both the Robofish and a live companion (N = 30). To further randomize our testing procedure, we performed Robofish and model fish trials in an alternating order at each experimental day.

All video recordings were subjected to a custom-made software [52] to extract position and orientation of both interaction partners over time. Based on the tracked positions, we calculated several measures that characterize social interactions (see [19, 54-58]).

As a simple proxy for the social interaction among subjects, we calculated the inter-individual distance (IID) between focal fish and companion (Robofish or live fish, body centroids) for each trial [2]. It is strongly correlated with other distance-related measures, such as the time fish spent within a specific range (not shown), and short distances between subjects suggest strong interactions (e.g., at least one subject must follow the companion closely).

As our major goal was to determine focal individual’s responsiveness towards its companions (Robofish or live model), we calculated subject-specific interaction measures for each individual (focal, live model as well as Robofish) within a pair. Freely interacting live fish respond rapidly to conspecifics’ movements by adjusting their own movement patterns [19, 54, 57-60]. To quantify this response in movement patterns, we calculated time-lagged cross-correlations of velocity vectors (TLXC), which allow to distinguish how strongly subject adjust their own movement towards that of the partner’s movement [61]. For any given time lag τ, TLXC indicates the strength of the correlation between the velocity vector of the focal individual at time t+τ and the other companion individual at time t. A large positive value implies that on average the focal individual responds by moving in the same direction as its companion, whereas values close to zero correspond to a random response and negative values indicate a movement in opposite direction. In a representative sample of our dataset, all first extrema in the cross-correlation can be found for lags < 6 s. We thus restricted our analysis to lag-times up to τ = 6 s. We calculated the cross-correlation averaged over the entire time lag window for both subjects within a pair. Subject-specific TLXCs were then used to calculate a global correlation measure as the difference between focal fish’s average cross-correlation and companion’s average cross-correlation (ΔTLXC; positive values: focal fish followed on average; negative values: focal fish led on average). When interactions are strong (e.g., when individuals are in close range), ΔTLXC values around zero indicate rapidly switching leadership roles/high mutuality in social responsiveness. When interactions are weak (e.g., at longer distances between fish), ΔTLXC values around zero indicate weak overall social responsiveness (for more details on the calculation of TLXC please see our Supplemental Information S1_Text1).

Transfer Entropy (TE) between velocity vectors of both individuals [62, 63]. TE_i⟶j_ is an information theoretic, model-free measure of directed (‘causal’) coupling between two time series *i* and *j*, and was shown to generalize Granger Causality to arbitrary nonlinear, stochastic processes [64]. For simplicity, we will use the short notation TE_i_ for the average TE_i⟶j_(τ), the Transfer Entropy between individual i and j at time lag τ. As for TLXC, we measured the average TE_i_ by averaging TE_i⟶j_(τ) over the entire time window (see SI_Text1) to quantify responsiveness. As the baseline values of TE_i_ depend on various factors not related to actual couplings, as e.g. activity of individuals, trajectory length and parameters of the used entropy estimator [65], we calculated the difference between focal fish’s average TE_i_ and companion’s average TE_j_ (ΔTE=TE_j⟶i_ (τ)-TE_i⟶j_ (τ)), which may be interpreted as the global net information flow with respect to velocity dynamics (see S1_Text1) between both individuals with the sign indicating the direction of the information flow and the absolute value indicating its strength. Positive values indicate that information is predominantly transferred from the companion (j) to the focal fish (i) while negative ones indicate the opposite. Values around zero suggest equal information transfer between subjects (when interactions are strong) or no information transfer (when interactions are weak).

##### Statistical analysis

In order to see whether the magnitude of social interactions between live pairs and Robofish pairs differed on average, we compared inter-individual distance (log-transformed), velocity cross-correlations (average TLXC of a pair as well as ΔTLXC) and Transfer Entropy (average TE of a pair as well as ΔTE) between live pairs and Robofish pairs using paired *t*-tests. In a second analysis, we compared TLXC and TE between subjects within Robofish as well as live fish pairs using paired samples *t*-tests. A correlational analysis between IIDs and TLXCs as well as TEs can be found as supplemental information (SI_Text2).

In our third analysis, we asked whether focal fish’s individual differences in responsiveness towards Robofish are mirrored in their interactions with live companions. We thus used univariate Linear Mixed Models (LMMs) with IID, TLXC (subject-specific and ΔTLXC)), and TE (subject-specific and ΔTE) as dependent variables and included focal fish ID as a random factor to calculate the behavioral ‘repeatability’ [49]. The repeatability of a behavior is defined as the proportion of the total behavioral variance (sum of variation that is attributable to differences among individuals plus variation within individuals) towards the amount of variation that is attributable to differences among individuals. As variance estimates are inherently tied to the total variation present in the response variable, we first mean-centered and scaled the variance of our response variables to 1 within each treatment (e.g., z-transformation). No fixed factors were included in the LMM to obtain conservative measures of among-and within-individual variation [49]. A significant repeatability estimate is interpreted as evidence of consistent individual differences and we tested for significance using likelihood ratio tests (see [66]).

#### Experiment 2

##### Experimental setup

The aim of our second experiment was to investigate a potential link between social responsiveness and other already established personality traits. Here, we were also interested in possible differences among the sexes and thus included males in our tests. To do so, male (N=17, SL = 19.5 mm ± 0.4 mm SEM) and female guppies (N=25, SL=27.6 ± 0.6 mm) were VIE tagged as described for experiment 1 and kept in 100-L tanks. After one week of acclimatization, all fish were tested three times for their personality types including their tendency to respond to Robofish.

##### Personality tests with Robofish

A test trial consisted of three consecutive parts. In the first part, focal fish were randomly taken from the stock tank and introduced into an opaque plastic cylinder with a small opening. The opening was closed with a sponge and fish were given 1 minute for habituation. Then, the sponge was removed and we scored the time each fish took to leave the cylinder as a measure of boldness (smaller values indicate bolder personalities, see [67, 68]). Robofish was positioned close to the opening at the outside of the cylinder so that the live fish could not see the Robot from the inside but could not miss it once it left the cylinder. Once the focal fish has left the cylinder, Robofish initiated the same zigzag sequence as described for experiment 1. However, this time Robofish did not move in a circular path, but was removed immediately after reaching one corner. To test for focal fish’s tendency to respond towards Robofish, we calculated subject-specific as well as pair-wise parameters described for experiment 1 as response variables. After Robofish was removed, focal fish were left in the tank and given 2 minutes for habituation. All fish resumed normal swimming within this 2 minutes and we videotaped them for another 3 minutes to get a measure of general activity (mean velocity,[69, 70]). Hence, each trial scored three different behavioral traits for which guppies are assumed to differ consistently among each other. Video analysis and parameter calculation followed the description provided for experiment 1.

##### Statistical analysis

To quantify how repeated testing or differences in sex and body size of the fish affected average behavioral traits, we analyzed ‘boldness’ (emergence time in s; log transformed prior to all statistical analysis), ‘social responsiveness’ (log-transformed IID, ΔTLXC and ΔTE; see experiment 1) and ‘general activity’ (mean velocity in cm/s) as dependent variables in separate LMMs with trial (three repeated test runs) and sex as fixed factors and focal fish’s body size (SL) as a covariate. Focal ID was included as random factor to account for repeated tests.

To see whether focal fish differed consistently in any of the behavioral traits, we used another set of LMMs with behavioral traits as dependent variables and Focal ID as a random factor. Similar to the analysis described for our first experiment, we first mean-centered and scaled the variance of our response variables to 1 within each trial (e.g., z-transformation).

We further asked whether our recorded behavioral traits may form a larger behavioral syndrome. We thus used Principle Component Analysis (PCA) with average values (z-transformed values averaged over the three trials) of pair-wise parameters (log emergence time, mean velocity, log IID, ΔTLXC and ΔTE) and extracted all components with Eigenvalues > 1. The resulting component matrix was Varimax rotated to ease interpretation of component loadings. Only loadings with values ≥ 0.8 were considered as important.

## Results

### Experiment 1

#### General response in pairs involving Robofish and only live fish

On average, general coupling between subjects was weaker in Robofish pairs than in live fish pairs. Distance between subjects was longer (paired *t*-test, IID: *t*_29_=-2.353; *P*=0.022, figure 3a), velocity correlations were less pronounced (average TLXC of both subjects: *t*_29_=-3.434; *P*=0.002; figure 3b) and information transfer rates were lower (average TE of both subjects: *t*_29_=-3.434; *P*=0.002; figure 3c) when focal fish were paired with Robofish as compared to a live companion.

**Figure 3:**
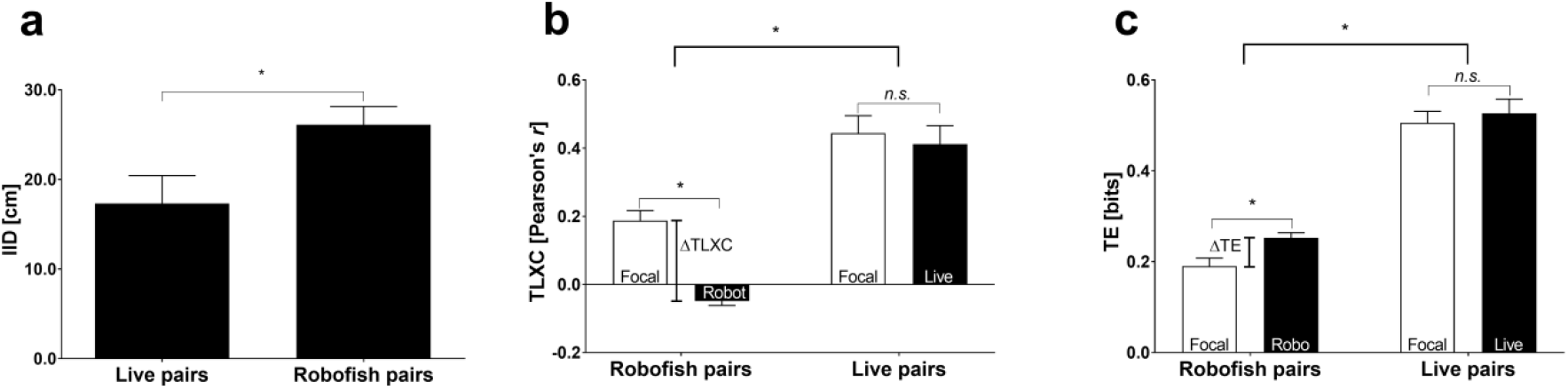
Average differences between Robofish pairs and live fish pairs in inter-individual distances (IID, a), time-lagged cross-correlation of velocity vectors (TLXC, b) and Transfer Entropy (TE, c). Shown are means ± SEM. Asterisks indicate significant differences in *t*-tests (see main text).

Robofish’s velocity vectors were not correlated with those of live focal fish as indicated by velocity vector cross-correlations (TLXC) of Robofish around zero that were significantly lower than those of the focal fish in Robofish pairs (*t*_29_=-6.613; *P*<0.001; figure 3b). In live pairs, both fish adjusted their velocities towards each other as indicated by high TLXCs that did not differ between subjects (*t*_29_=-0.901; *P*=0.375; figure 3b). As a result, ΔTLXC was significantly higher in Robofish pairs compared to live fish pairs (*t*_29_=-4.031; *P*<0.001; figure 3b).

Regarding TE, the lack of response of the Robofish led to a significant difference in TE values between live focal fish and Robofish (*t*_29_=-5.442; *P*<0.001; figure 3c) with a net information flow towards focal fish (positive ΔTEs, figure 3c). This pattern was not seen in live pairs where TEs did not differ between subjects (*t*_29_=-1.259; *P*=0.218; figure 3c) as information flow was shared equally (small ΔTEs, significantly different to that found in Robofish pairs; *t*_29_=-2.001; *P*=0.043, figure 3c).

Overall, our results indicate that focal fish in Robofish pairs were predominately adjusting their own swimming behavior to that of Robofish and not vice versa, while focal fish and live model companions within live pairs were mutually responding towards each other. Thus, the general difference in interaction patterns between Robofish pairs and live fish pairs are likely due to the mutual responses that we see in live fish pairs but, as intended, not in Robofish pairs.

#### Individual differences in social responsiveness

We hypothesized that a live fish’s reaction towards Robofish should reflect its social responsiveness, similar as to when tested with another live companion. Although there are general differences among response towards Robofish and a live companion, we found that focal individuals differed consistently across treatments with regard to all subject-specific interaction parameters as well as inter-individual distances (IID) and Robofish’s TE and ΔTE (table 1a). Only companions’ TLXC and ΔTLXC were not repeatable. This is most probably due to the discrepancy between high TLXC values in live companions (highly responsive) and the low TLXC values of Robofish (non-responsive; see figure 3b). Interestingly, focal individuals’ influence on companions’ TE and ΔTE seem to be strong enough to even detect consistent individual differences here.

**Table 1:**
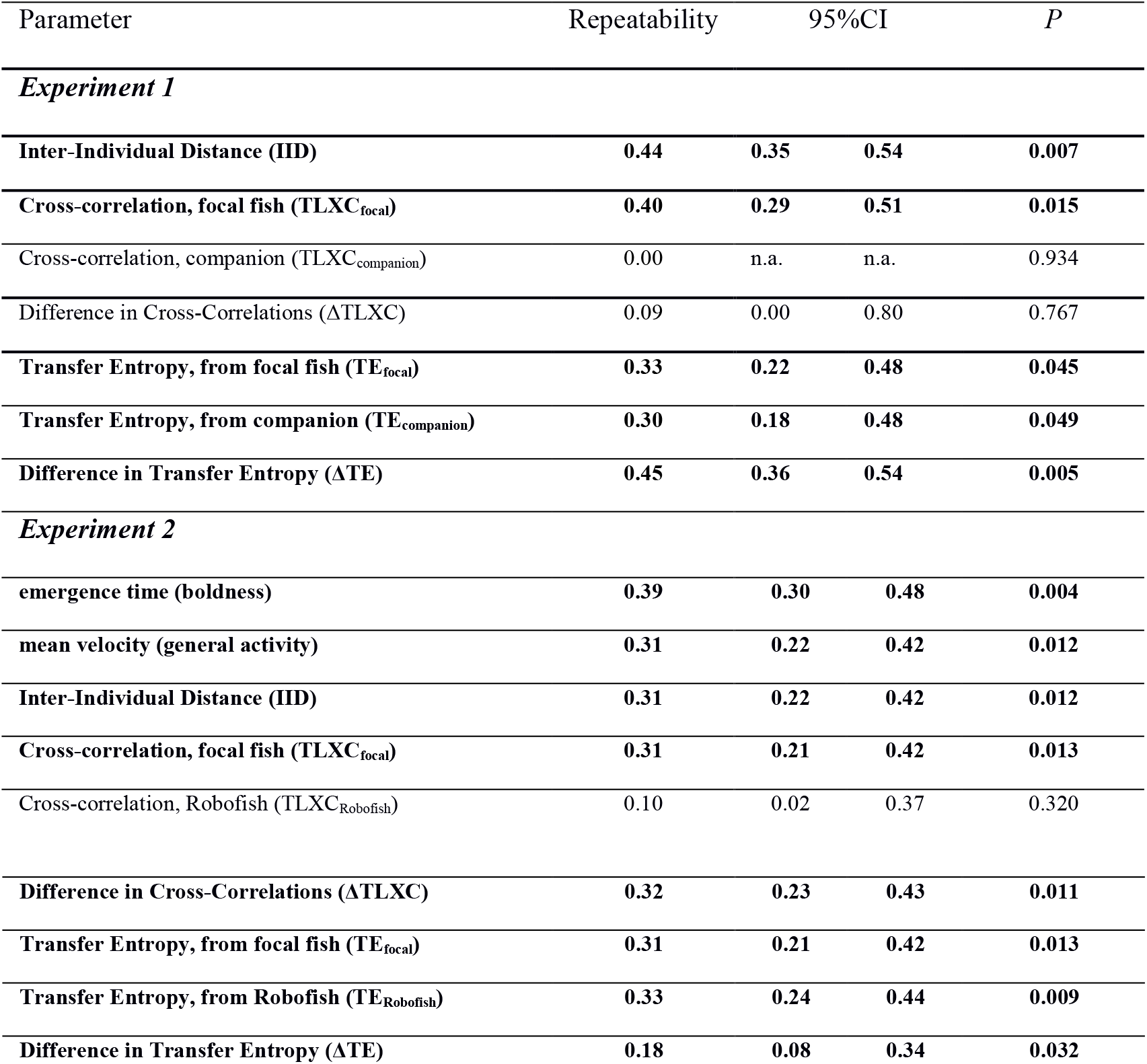
Behavioral repeatability of subject-specific and pair-wise interaction parameters. Shown are repeatability values obtained from LMMs on treatment-centered and normalized parameters along with 95% credibility intervals and significance levels from likelihood ratio tests. A significant repeatability indicates consistent individual differences. Significant repeatability values are in bold type face.

### Experiment 2

#### Is Social Responsiveness part of a larger behavioral syndrome?

On average, boldness scores were not affected by repeated testing, i.e. there was no effect of habituation (*F*_2,82_=0.092, *P*=0.911; figure 4a). However, focal fish decreased their general activity significantly over time (sig. effect of factor ‘trial’ in LMM: *F*_2,82_=9.111, *P*<0.001, figure 4b). A similar effect was observed for social responsiveness parameters (IID: *F*_2,82_=30.908, *P*<0.001, figure 4c; ΔTLXC: *F*_2,82_=11.737, *P*<0.001, figure 4d; ΔTE: *F*_2,82_=5.683, *P*=0.005, figure 4e) and focal fish adjusted their velocities to a lesser extent and kept longer distances towards Robofish with repeated testing. Interestingly, net information flow towards focal fish (ΔTE) increased with repeated testing. Body length of the test fish had no significant effect in either model (not shown), while males received more information from Robofish than females (ΔTE: sig. effect of factor ‘sex’: *F*_1,39_=9.626, *P*=0.004).

**Figure 4:**
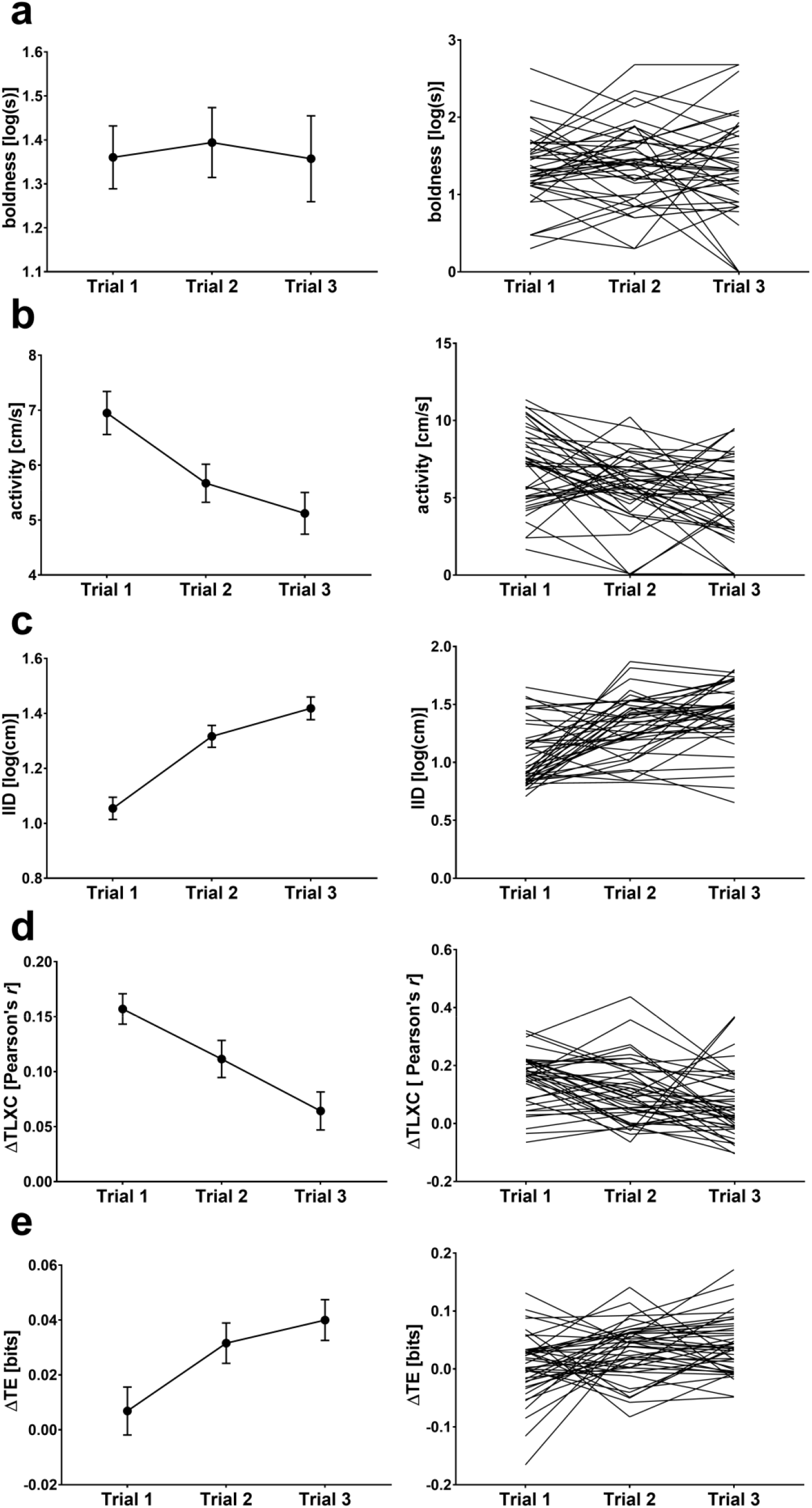
Behavioral traits in the guppy. (left) Average values for behavioral traits across the repeated testing. Shown are means ± SEM. Please note that except for ‘boldness’, trial 1 was always significantly different from trial 2 and 3 (*post-hoc* LSD tests). (right) Individual values for focal fish. Each line shows a focal individual’s behavioral expression over the repeated testing.

Our repeatability analysis found that focal fish differed consistently in all three behavioral traits (Table 1, figure 4, right side). This repeatability indicates that focal fish largely maintained their individual differences in boldness and activity as well as responsiveness when interacting with Robofish over the course of the repeated testing.

Principle component analysis (PCA) found 3 components with Eigenvalues >1 that explained 84.6% of the total variance in the data. Boldness and activity loaded strongest on component 2 (PC2, 24.5% variance explained; figure 5), which suggests that bolder fish (i.e., those that left the start cylinder quicker) were also more active (negative relation). IID and ΔTLXC loaded strongest on PC1 (39.5% variance explained), and, similar to the results from experiment 1, ΔTLXCs were high in pairs where focal fish were also close to Robofish (see SI_Text2) while ΔTLXCs decreased when IIDs increased (different signs in component loadings, see figure 5). ΔTE loaded independent of other variables on PC3 (20.6% variance explained). To sum up, our proxies for social responsiveness (IID, ΔTLXC, ΔTE), although repeatable, were not correlated with boldness or activity and, thus, do not seem to be part of a larger boldness-activity syndrome.

**Figure 5:**
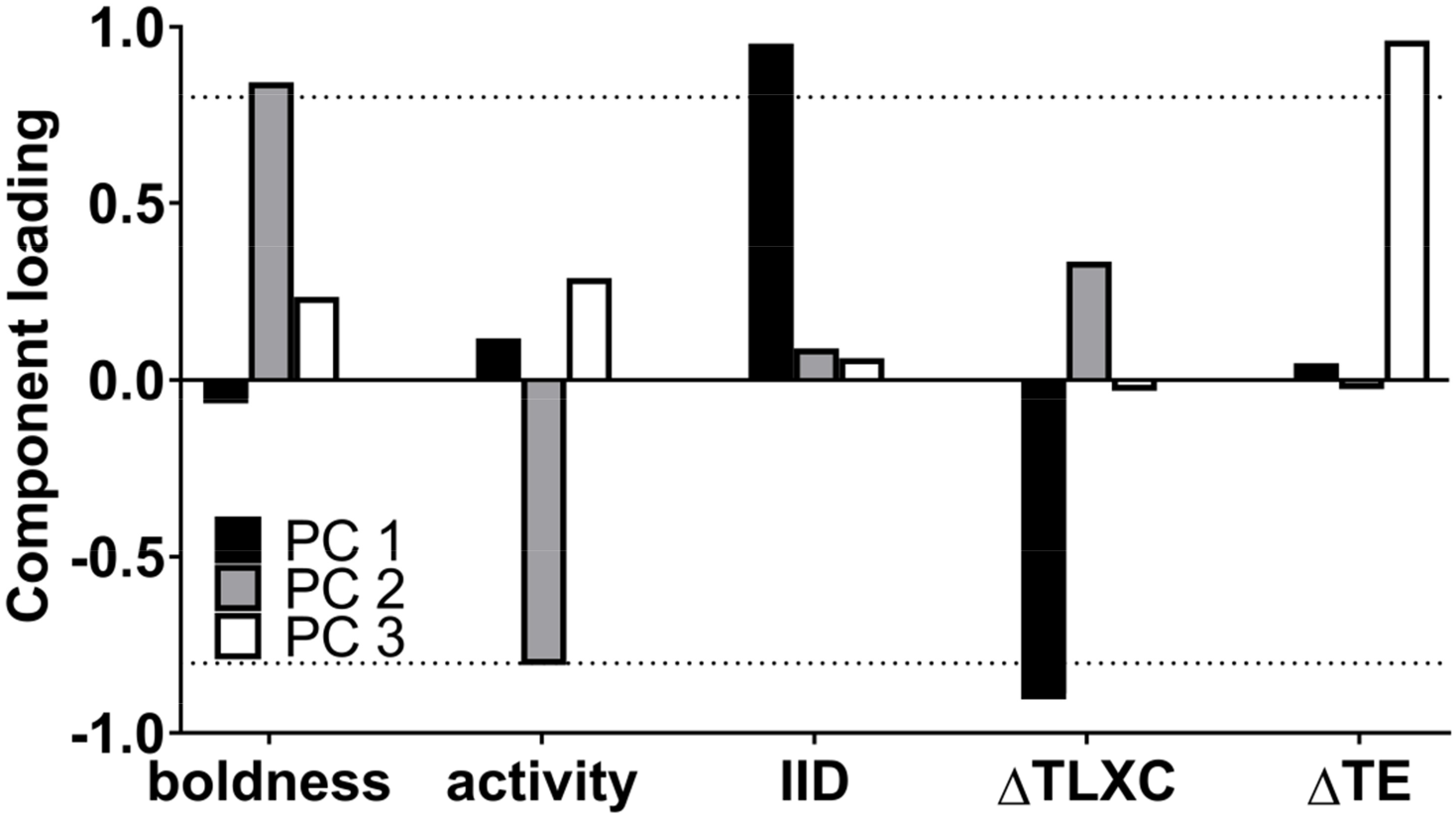
Results from Principle Component Analysis (PCA) on pair-wise behavioral traits. Shown are component loadings for the first three PCs with Eigenvalues >1. Dashed line at 0.8 indicates important loadings.

## Discussion

Our results show that guppies were consistent in their individual responses between a live companion and a robotic companion - the Robofish (first experiment). Furthermore, these individual differences are maintained over repeated testing with Robofish even though habituation to the test tank is detectable (second experiment). In addition, guppies differed consistently in their boldness and general activity with a similar habituation to the test tank found for the latter. While our boldness and activity measures were correlated (e.g., seem to form a behavioral syndrome with bolder individuals being also more active), social responsiveness towards Robofish was not correlated with boldness and activity measures and thus independent of the boldness-activity behavioral syndrome.

In our first experiment, we aimed to validate our approach as biologically meaningful (see [71]) by showing that interactions of live fish with our robot (a) contain elements that are typical of pair-wise interactions in live fish and (b) that they reflect individual differences in guppies’ social responsiveness. Indeed, guppies readily followed the moving Robofish and mean distances between subjects largely overlapped with those recorded for live fish pairs, although live fish were on average closer to each other. In regard to our measures of social responsiveness (velocity cross-correlations [TLXC] and transfer entropy [TE]), focal guppies tested with Robofish showed comparable albeit weaker response patterns as when tested with live companions. Nevertheless, live guppies paired with Robofish adjusted their velocities in accordance to Robofish (positive TLXC of live fish in Robofish pairs) especially when in close range (SI_Text2). Similarly, our measure of information transfer (TE) found live guppies to integrate directional information received from Robofish into their own movement patterns (information transfer from Robot to live fish) which was greatest when subjects were close together (SI_text2).

The major difference between interactions of live fish pairs and pairs involving Robofish was the strikingly high mutual responsiveness of individuals in live pairs (no significant differences in TLXC and TE within live pairs). In contrast, Robofish’s (intended) unresponsiveness was well detectable (zero TLXC, low information flow from live fish to Robofish) leading to significant differences in net (Δ) TLXC and net (Δ)TE compared to live fish pairs. These differences may have further caused the greater inter-individual distances among subjects in Robofish pairs. Despite these differences, focal fish maintained individual differences (sig. repeatability) in individual interaction parameters across both test situations (i.e., with a live companion and a Robofish partner). We are, thus, confident that reactions towards Robofish provide a consistent and reliable measure for social responsiveness in live guppies (see similar validations for Zebrafish’s (*Danio rerio*) responses towards biomimetic robots [32] and experiments with sticklebacks and circulating replica shoals [42]).

While individual differences in behavior seem to be an inevitable outcome of complex development and arise even in the absence of genetic and/or environmental variation [70], many personality traits form larger correlated suits, so-called ‘behavioral syndromes’ [72]. In our second experiment, we show that guppies differed consistently in boldness, activity, as well as social responsiveness towards Robofish. However, responsiveness towards Robofish was not correlated with boldness and activity and thus most likely not part of a larger behavioral syndrome formed around a boldness-activity axis. The correlation between boldness and activity is well described for poeciliid fishes including the guppy [69, 73-76]. As a point of caution, we note that we might overestimate correlations between boldness and activity as we measured both traits in short succession at the same day (see [77] for a discussion).

In many species, bolder individuals are more likely to initiate exploration of new environments and thus are assumed to have a greater tendency to lead others [8, 15, 21, 78-82]. In sticklebacks, bolder individuals are often less socially attracted (‘less sociable’) and less responsive to their current social partners [8, 23]. Both traits are assumed to result in lower tendencies to follow other group members [15], yet this does not hold true for our results as responsiveness towards Robofish was not correlated with boldness. We do not have a compelling explanation for this discrepancy but might argue that our study design did not allow any kind of mutual feedback, which is known to modulate leader-follower interactions [81, 83] and is an inevitable property of any tests involving multiple live animals [36]. One possibility to investigate the effect of mutual but still controllable feedback would be to use interactive robots (closed-loop mode) that respond to live subjects [45, 48].

## Acknowledgments

We would like to thank Nadine Muggelberg, Romain J.G. Clément, Joseph Schröer, Angelika Szengel, Marie Habedank, Hauke J. Mönck as well as Hartmut Höft and David Lewis for their valuable help in the lab.

## Ethics

The experiments reported herein comply with the current German laws approved by LaGeSo Berlin (Reg 0117/16 to D.B.).

## Author contributions

D.B., T.L., M.W. and J.K. conceived the study; D.B., J.L. and H.N. performed the experiments; D.B. and P.R. analyzed the data; D.B., T.L. and H.N. developed the Robofish system; D.B. wrote the manuscript. All authors contributed to and approved the submitted version of the manuscript.

## Funding

This research was financed by the DFG (to D.B.: BI 1828/2-1; to P.R.: RO 4766/2-1; to T.L.: LA 3534/1-1) as well as the Leibniz Competition (B-Types project; SAW-2013-IGB-2 to M.W. and J.K.).

## Competing interests

The authors declare no competing interests.

## Supplemental Information

**Using a robotic fish to investigate individual differences in social responsiveness in the guppy**

David Bierbach^*^, Tim Landgraf^*^, Pawel Romanczuk, Juliane Lukas, Hai Nguyen, Max Wolf, Jens Krause

^*^ Authors contributed equally

### SI_TEXT1: Calculation of interaction parameters

#### Average time-delayed cross-correlation (TLXC)

We calculated first the time-delayed normalized cross correlation function for different values of the lagtime *τ*:*C(τ)=< v(t)v_f_(t+τ)>_t_* with *v_f_* being the velocity of the focal individual and <…>_*t*_ indicating a time average. Then we calculated **TLXC** as the average *C(τ)* over a finite range of timelags (0-6s): **TLXC**=< *C(τ) >*

#### Average Transfer Entropy (TE)

The transfer entropy is calculated based on velocity vectors as input variables. First we calculate the transfer entropy for a given timelag using:

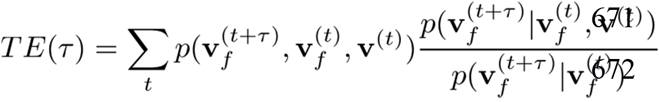

with v_f_^(t+τ)^ and v_f_^(t)^ being the velocity vectors of the focal individual at time t+ τ (future) and t (past), respectively, and v^(t)^ being the velocity of the other individual at time t. Here, *p*(**v**_*f*_^(*t*+ τ)^, **v**_*f*_^(*t*)^, **v**^(*t*)^) is the overall probability of observing the corresponding values of the three velocity vectors, *p*(**v**_*f*_^(*t*+ τ)^|**v**_*f*_^(*t*)^, **v**^(*t*)^) is the probability to observe future velocity vector of the focal individual v_f_^(t+τ)^ conditioned on the past velocity of the focal individual as well as the other individual, whereas *p*(**v**_*f*_^(*t*+ τ)^|**v**_*f*_^(*t*)^) is the probability to observe v_f_^(t+τ)^ conditioned only on the past velocity of the focal individual. For efficient estimation of the transfer entropy we used the k-nearest neighbor estimator [1-3]. The averaged TE was then obtained by averaging the TE(τ) over a range of lagtimes τ (0-6s).

## SI_TEXT2: Additional results for Experiment 1

### Measures of social responsiveness: Velocity vector cross-correlations (TLXC) and inter-individual distances (IID)

We found that focal fish’s (albeit weaker) responses to Robofish resembled those shown towards live conspecifics: Focal fish adjusted their movement patterns (velocity vectors) rapidly to those of their respective partners (both live companions and Robofish) when in close range and this response became weaker with increasing distance among subjects (figure S1a,b). This was indicated by a significantly negative correlation between inter-individual distance (IID) and focal fish’s time-lagged velocity vector cross-correlation (TLXC) both in Robofish pairs (focal fish: *r*_pearson_=-0.82; *N*=30, *P*< 0.001; figure S1a) as well as in live fish pairs (focal: *r*_pearson_=-0.79; *N*=30, *P*< 0.001; figure S1b).

Regarding the companions, live model fish responded with a similar adjustment of own velocity vectors towards the focal fish and cross-correlations decreased similarly with increasing distance among subjects (companion: *r*_pearson_ =-0.68; *N*=30, *P*<0.001; figure S1b). However, the non-interactive Robofish did not adjusted its movement towards the focal fish at any distance (no correlation detectable for Robofish’s TLXC; *r*_pearson_=0.2; *N*=30, *P*=0.32; figure S1a).

**Figure S1:**
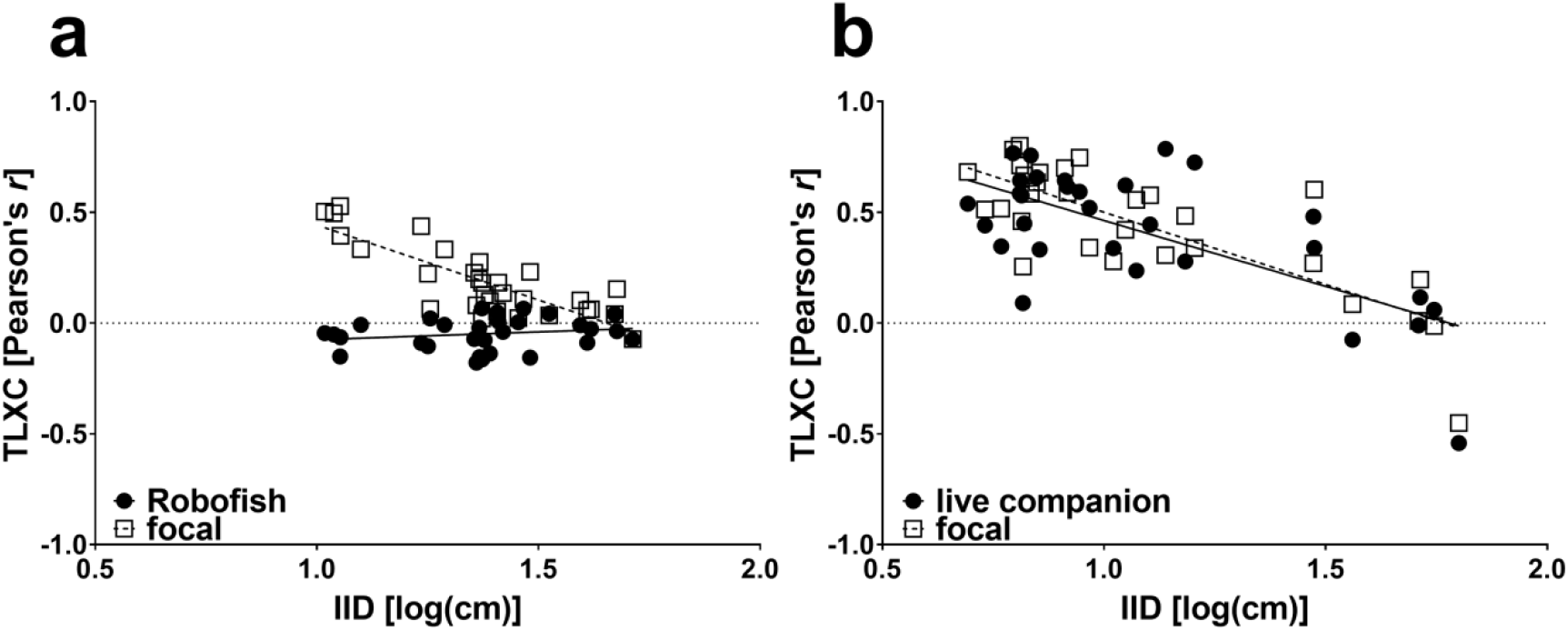
Time-lagged cross-correlation of individual velocity vectors (TLXC) and inter-individual distances (IID). (a) In Robofish pairs, only live fish’s TLXCs were correlated with inter-individual distance (IID) but not Robofish’s TLXC. (b) In live fish pairs, both subjects showed a correlation between TLXC and IID.

### Measures of social responsiveness: Transfer Entropy (TE) and inter-individual distance (IID)

As for velocity cross-correlations, levels of information transfer towards focal fish in Robofish pairs became higher the closer the subjects were (sig. negative correlation between IID and Robofish’s TE in Robofish pairs; *r*_pearson_=-0.584; *N*=30, *P*<0.001; figure S2a), while the TE from live fish to Robofish did not correlate with distance between subjects (no correlation detectable for live fish’s TE_i_ with IID in Robofish pairs; *r*_pearson_=-0.350; *N*=30, *P*=0.058; figure S2a).

Interestingly, TE for both subjects in live pairs was independent of inter-individual distance (IID) (no correlations; focal: *r*_pearson_=0.08; *N*=30, *P*=0.658; companion: *r*_pearson_=0.02; *N*=30, *P*=0.91; figure S2b).

**Figure S2:**
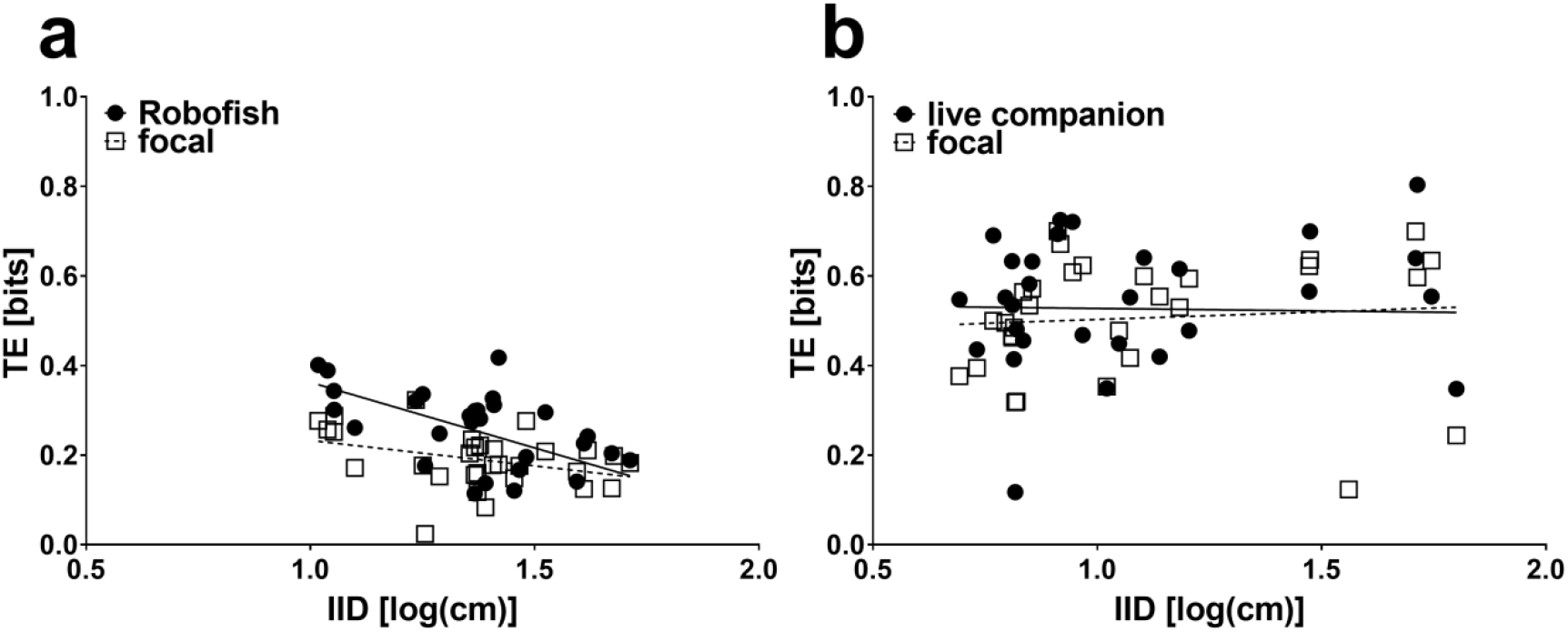
Transfer Entropy (TE) and inter-individual distances (IID). (a) In Robofish pairs, only Robofish’s TE were correlated with inter-individual distance (IID) but not live fish’s TE. (b) In live fish pairs, both subjects showed no correlation between TE and IID.

